# A Nanoparticle RIG-I Agonist for Cancer Immunotherapy

**DOI:** 10.1101/2023.04.25.537919

**Authors:** Lihong Wang-Bishop, Mohamed Wehbe, Lucinda E. Pastora, Jinming Yang, Kyle M. Garland, Kyle W. Becker, Carcia S. Carson, Katherine N. Gibson-Corley, David Ulkoski, Venkata Krishnamurthy, Olga Fedorova, Ann Richmond, Anna Marie Pyle, John T. Wilson

## Abstract

Pharmacological activation of the retinoic acid-inducible gene I (RIG-I) pathway holds promise for increasing tumor immunogenicity and improving response to immune checkpoint inhibitors (ICI). However, the potency and clinical efficacy of 5’-triphosphate RNA (3pRNA) agonists of RIG-I is hindered by multiple pharmacological barriers, including poor pharmacokinetics, nuclease degradation, and inefficient delivery to the cytosol where RIG-I is localized. Here, we address these challenges through the design and evaluation of ionizable lipid nanoparticles (LNPs) for the delivery of 3p-modified stem-loop RNAs (SLRs). Packaging of SLRs into LNPs (SLR-LNPs) yielded surface charge-neutral nanoparticles with a size of ∼100 nm that activated RIG-I signaling *in vitro* and *in vivo*. SLR-LNPs were safely administered to mice via both intratumoral and intravenous routes, resulting in RIG-I activation in the tumor microenvironment (TME) and inhibition of tumor growth in mouse models of poorly immunogenic melanoma and breast cancer. Significantly, we found that systemic administration of SLR-LNPs reprogrammed the breast TME to enhance the infiltration of CD8^+^ and CD4^+^ T cells with antitumor function, resulting in enhanced response to αPD-1 ICI in an orthotopic EO771 model of triple negative breast cancer. Therapeutic efficacy was further demonstrated in a metastatic B16.F10 melanoma model, with systemically administered SLR-LNPs significantly reducing lung metastatic burden compared to combined αPD-1 + αCTLA-4 ICI. Collectively, these studies have established SLR-LNPs as a translationally promising immunotherapeutic nanomedicine for potent and selective activation of RIG-I with potential to enhance response to ICIs and other immunotherapeutic modalities.

## Introduction

Immunotherapy has revolutionized the treatment of an expanding diversity of tumor types, resulting in robust and durable responses for some patients.^1^ However, it is now well-recognized that, for most cancer types, only a minority of patients respond to currently approved immune checkpoint inhibitors (ICI) that target CTLA-4 and PD-1/PD-L1.^2^ While resistance to ICIs is multifaceted, for many cancer types the response to ICI correlates with an immunogenic (“hot”) tumor microenvironment (TME) that is infiltrated with tumor antigen-specific CD8^+^ cytotoxic T cells that are reactivated by immune checkpoint blockade (ICB).^2, 3^ However, accumulating data indicate that many patients – perhaps the majority – have immunologically “cold” tumors that lack significant T cell infiltration and are instead characterized by a high density of immunosuppressive cells that inhibit antitumor immunity. This has motivated wide-spread investigation into the development of therapeutics that reprogram the TME to increase the number and/or function of tumor infiltrating T cells that can be reactivated in response to ICIs.^3–5^

Innate immunity plays a critical role in the detection and elimination of cancers.^6^ The innate immune system employs pattern recognition receptors (PRRs) – a network of molecular sensors that detect distinctive features of pathogens or damaged tissue (i.e., ‘danger signals’) – to trigger inflammatory responses that are critical to the recruitment of immune cell populations to sites of infection, tissue injury, and malignancy.^6, 7^ Retinoic acid-inducible gene-I (RIG-I) (also known as DDX58) is an important PRR for sensing of RNA viruses^8^ via recognition of short, double-stranded RNA with a triphosphate group (3p) on the 5’ end (3pRNA).^9, 10^ The 3p group acts as a ‘tag’ that allows RIG-I to discriminate between 3pRNA and other cytosolic RNAs (e.g., mRNA, miRNA, etc.) with high selectivity. Activation of RIG-I triggers a multifaceted innate immune response characterized by the expression of type-I interferons (IFN-I), interferon stimulated genes (ISGs), T cell chemokines (e.g., CXCL-9, 10), and proinflammatory cytokines, which cooperate to exert direct and broad-spectrum antiviral functions while also augmenting and shaping the subsequent adaptive immune response.^11–13^ Evidence is also emerging that RIG-I can detect self RNA derived from aberrantly expressed endogenous retroviral elements dispersed within the human genome or mislocalized mitochondrial RNA in the cytosol and, hence, may also have an important role in promoting endogenous immunity against cancer.^14, 15^ Indeed, RIG-I signaling in cancer cells has recently been shown to mediate responsiveness to anti-CTLA-4 immune checkpoint blockade in tumor-bearing mice, consistent with an association between RIG-I expression level, T cell infiltration, and overall survival in melanoma patients.^16^ Such links between RIG-I and endogenous cancer immune surveillance motivates the development of RIG-I agonists as cancer immunotherapies.

While promising as an immunotherapy agent, 3pRNA RIG-I agonists face multiple barriers to therapeutic efficacy that are shared with many oligonucleotide therapies, including a short plasma half-life (i.e., minutes), high susceptibility to nuclease degradation, poor intracellular uptake, and, critically, degradation in endo/lysosomes with minimal delivery to the cytosol where RIG-I is localized.^17, 18^ In considering this drug delivery challenge, we postulated that clinically-advanced lipid nanoparticle (LNP) technology could be harnessed to overcome barriers to 3pRNA delivery, thereby opening a therapeutic window for pharmacological activation of RIG-I as a cancer immunotherapy. LNPs have been widely employed for the delivery of diverse types of nucleic acid therapeutics (e.g., mRNA, siRNA, DNA).^19, 20^ Their capacity to efficiently package and facilitate the cytosolic delivery of drug cargo is vital to the success of several FDA-approved LNP-based nanomedicines, including the Moderna and Pfizer-BioNTech mRNA COVID-19 vaccines.^21^ However, LNP formulations of 3pRNA have not yet been explored for immunotherapeutic activation of RIG-I.

Here, we describe the design and pre-clinical evaluation of a nanoparticle RIG-I agonist for cancer immunotherapy based on a remarkably simple yet highly effective approach. We leveraged an ionizable lipid that is already used in an FDA-approved siRNA therapeutic^22^ to package a 3p-modified, stem-loop RNA (SLRs) that we have engineered to be a molecularly-defined, selective, and high-affinity RIG-I agonist.^11^ We found that SLR-loaded LNPs (SLR-LNP) inhibited tumor growth and increased response to ICIs in poorly immunogenic, orthotopic mouse models of breast cancer and melanoma. Importantly, whereas most previous reports^13, 23, 24^ and early-stage clinical trials (e.g. NCT03739138)^25^ have relied on intratumoral injection of 3pRNA complexed to the cationic transfection agent JetPEI^TM^, SLR-LNPs could be safely administered systemically via intravenous injection, resulting a nearly complete elimination of lung metastases in mice with ICI-resistant metastatic melanoma. Collectively, our studies have yielded amongst the most potent and effective strategies for pharmacological RIG-I activation described to date and have identified LNPs as a previously unexplored and translationally-ready nanotechnology platform for harnessing the potential of RIG-I in cancer immunotherapy.

## Results

### Lipid nanoparticle delivery of SLR potently activates RIG-I signaling

LNPs are comprised of several types of lipids, including ‘helper’ lipids that contribute to structure and delivery efficiency, lipids modified with poly(ethylene glycol) (PEG-lipid) to confer colloidal stability and blood compatibility, and, importantly, ionizable lipids that facilitate packaging of RNA cargo via electrostatic interactions and promote the delivery of RNA into the cytoplasm following endocytosis and endosomal acidification.^19, 20^ While an ever expanding number of novel ionizable lipids are being developed, few are currently approved as components of therapeutics that are administered systemically (i.e., intravenously) in humans.^26^ Therefore, we selected DLin-MC3-DMA (MC3), a component in the FDA-approved, siRNA therapeutic ONPATTRO (patisiran)^22, 26^ as a translationally-relevant ionizable lipid for our design (**Figure 1A**). To confer colloidal stability and improve circulation half-life, 3.5% PEGylated lipid (DMG-PEG2000) was used in the formulation as well as 7.5% cholesterol and 31.5% DSPC (1,2-distearoyl-sn-glycero-3-phosphocholine) as helper lipids. We increased the amount of PEGylated lipid relative to that used in the patisiran formulation (3.5% vs. 1.5%) based on previous work demonstrating that increased PEGylation can increase circulation time and enhance tumor accumulation.^27^ In our design we also employed a well-defined, high-affinity, stem-loop 3pRNA (SLR) ligand for RIG-I that we have previously leveraged for potent and specific pharmacological activation of RIG-I in mice.^11, 17^ SLRs are synthesized using solid phase nucleic acid synthesis methods, enabling high yield and purity of potent RIG-I agonists with advantages over double-stranded 3pRNA synthesized via *in vitro* transcription that have been primarily utilized. Here, we used SLR20, a single-stranded 44-mer that folds into a stem-loop structure with a 20 base pair stem and a four-nucleotide loop (**Figure 1B**).

**Figure 1.**
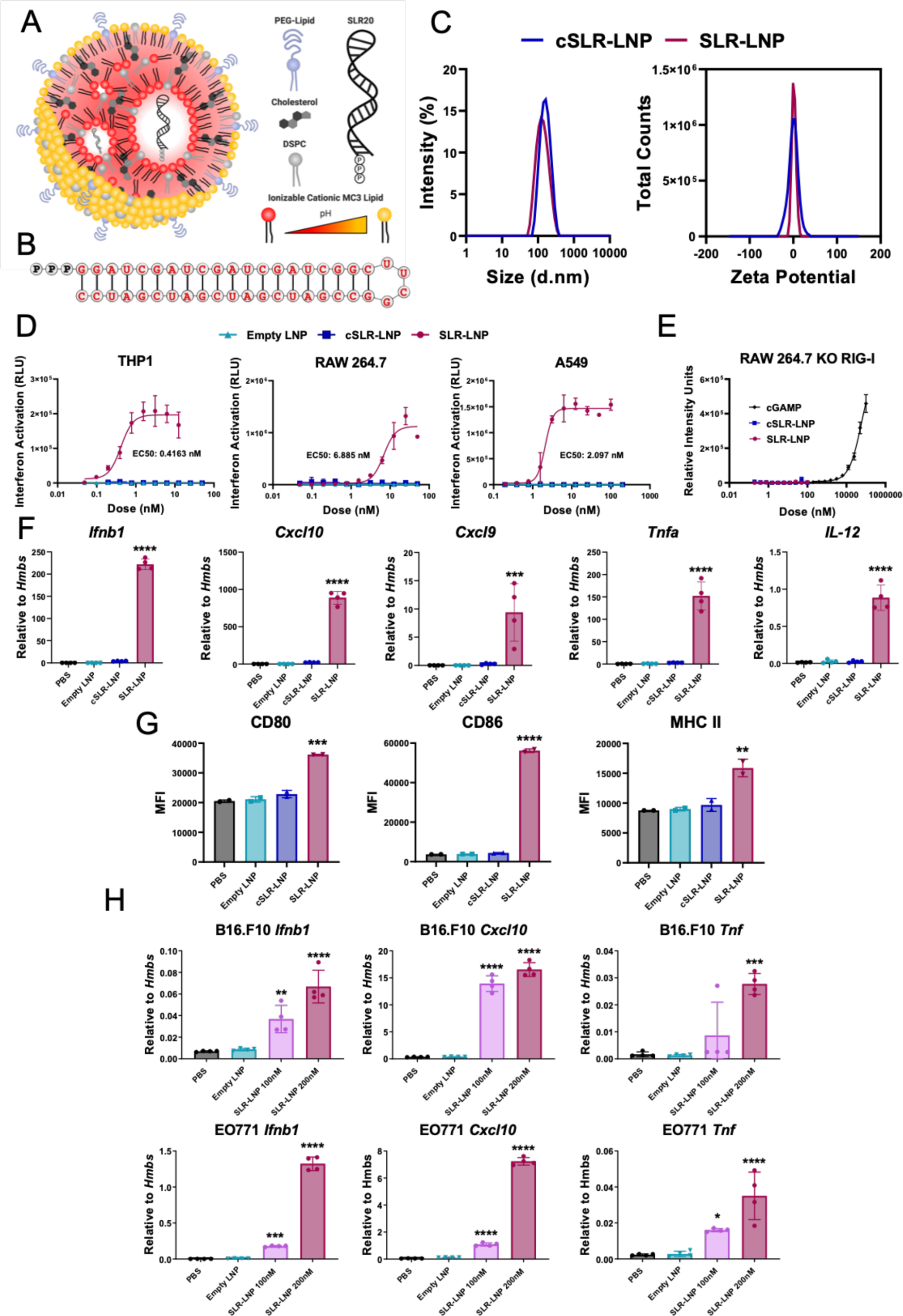
Lipid nanoparticle delivery of SLR20 potently activates RIG-I. **(A)** Schematic of SLR20-LNP composition and structure. **(B)** Structure and sequence of SLR20. **(C)** Particle size distribution determined by dynamic light scattering and zeta potential at pH 7.4 of LNPs loaded with SLR20 and SLROH. **(D)** Dose-response curves for type-I IFN (IFN-I) elicited by indicated LNP formulations in THP-1, RAW264.7, and A549 cells with an IFN regulatory factor (IRF)-inducible reporter construct. **(E)** Dose-response curve of type-I IFN (IFN-I) response elicited by indicated LNP formulation or STING agonist cGAMP (positive control) in RAW264.7 KO-RIG-I cells with an IFN regulatory factor (IRF)-inducible reporter construct. **(F)** RT-qPCR analysis of bone marrow derived dendritic cells (BMDCs) treated with indicated LNP formulation or PBS for 24h. (n=4, *** p<0.001, ****p<0.0001, by one-way ANOVA). **(G)** Flow cytometric quantification (median fluorescent intensity) of CD80, CD86, and MHC II expression on BMDCs stained after 24h treatment with indicated LNP formulation or PBS. (n=2, **p<0.01, *** p<0.001, ****p<0.0001, by one-way ANOVA). **(H)** RT-qPCR analysis of B16.F10 melanoma cells and EO771 breast cancer cells treated with indicated LNP formulation or PBS for 24h. (n=4, *p<0.05, **p<0.01, *** p<0.001, ****p<0.0001, by one-way ANOVA). All statistical data are presented as mean ± SD.

Mixing of lipids and SLR20 in citrate buffer (pH 3) at a nitrogen:phosphate (N:P) ratio of 4.8:1 resulted in the assembly of SLR-loaded LNPs (**SLR-LNP**) with near 100% RNA encapsulation efficiency, a diameter of ∼100 nm, and a neutral zeta potential (**Figure 1C**).The immunostimulatory activity of SLR-LNP was evaluated in a series of type-I interferon (IFN-I) reporter cell lines, with dose-response studies yielding EC_50_ values in the 1 to 10 nM range, depending on cell type (**Figure 1D**). Importantly, empty LNPs and LNPs loaded with an analogous negative control SLR (cSLR) that lacked the 3p moiety and instead displayed a 5’-hydroxyl group did not induce an IFN-I response. Using RIG-I knockout reporter cells, we also validated that the IFN-I response induced by SLR-LNPs was dependent on RIG-I (**Figure 1E**). We also tested the activity of SLR-LNPs in primary murine bone marrow-derived dendritic cells (BMDCs), finding that SLR-LNPs, but not empty LNP and cSLR-LNP controls, stimulated expression of IFN-I (*Ifnb1*), interferon-stimulated genes (ISGs) (*Cxcl9*, *Cxcl10*), and Th1 cytokines (*Tnfa*, *Il12*) (**Figure 1F**), and increased surface expression of the dendritic cell (DC) activation and maturation markers CD80, CD86, and MHC-II (**Figure 1G**). Finally, since we, and others, have demonstrated that RIG-I activation in cancer cells can be important to therapeutic responses,^16, 28, 29^ we also tested activity of SLR-LNPs in B16.F10 melanoma and EO771 breast cancer cells, again demonstrating that SLR-LNPs increased expression of cytokines associated with RIG-I activation relative to controls (**Figure 1H**). Hence, LNPs provide a facile strategy for efficient packaging and intracellular delivery of SLR20, yielding a new class of immunostimulatory nanoparticle with broad potential clinical utility.

### SLR-LNPs stimulate antitumor innate immunity

We evaluated the *in vivo* activity of SLR-LNPs in a poorly immunogenic B16.F10 melanoma model, first using the intratumoral administration route that has been most commonly employed for evaluation of RIG-I agonists,^16, 24^ including in recent clinical trials (**Figure 2**).^25^ Consistent with *in vitro* data, SLR-LNPs promoted expression of pro-inflammatory, antitumor cytokines (*Ifnb1*, *Tnfa*, and *Il12*) as well as *Cxcl9* and *Cxcl10*, important chemokines for directing T cell infiltration (**Figure 2A**). We also tested the immunostimulatory activity SLR-LNP in a melanoma model in which B16.F10 cells express an IFN-inducible luciferase reporter, demonstrating that intratumoral administration of SLR-LNPs increases IFN signaling in the tumor cell compartment (**Figure 2B**).

**Figure 2.**
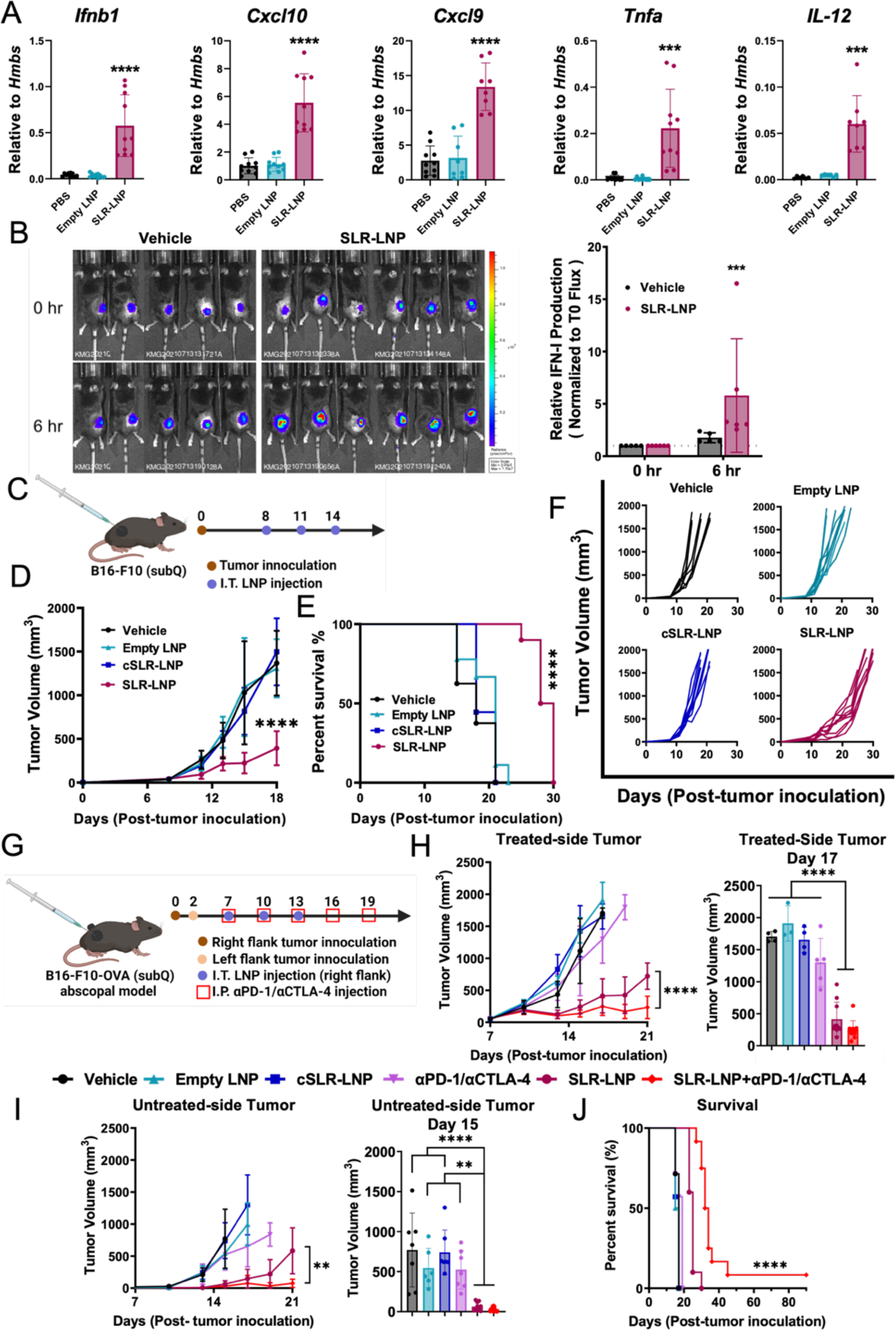
Intratumoral administration of SLR-LNPs activate RIG-I in the tumor microenvironment to inhibit local and distal tumor growth. **(A)** RT-qPCR analysis of B16.F10 tumors after intratumoral injection with single 100μL dose of either SLR-LNP (10 μg SLR), empty LNP, or PBS (n = 10, ***P ≤ 0.001; **** P ≤ 0.0001 by one-way ANOVA.) (B) *Left:* Representative luminescence IVIS images of IFN production in B16.F10 IFN-LUC tumors. An intratumoral injection was given as single 100 μL dose of either SLR-LNP (10 μg SLR) or PBS. *Right:* statistical analysis of relative IFN activity following treatment; data was normalized to t=0 h. (***P ≤ 0.001 by paired t-test, n=6). **(C)** Schematic representation of the B16-F10 melanoma single subcutaneous tumor model showing the treatment schedule. **(D)** Growth curves (**** P ≤ 0.0001 by one-way ANOVA compared to all other groups at day 18), **(E)** Kaplan-Meier survival curves, and **(F)** spider plots for B16-F10 single subcutaneous tumors. Tumors were measured by a caliper every other day from day 8 through 30 after tumor implantation (n=8-10 mice per group). **(G)** Schematic representation of the subcutaneous B16.F10-OVA melanoma abscopal tumor model showing the treatment schedule. **(H)** Growth curves of the treated-side tumors (H, *left*, **** P ≤ 0.0001 at day 21 by paired t-test), and volumes of treated tumors on day 17 (H, *right*, **** P ≤ 0.0001 by one-way ANOVA). **(I)** Growth curves of the untreated-side tumors (I, *left*, **P ≤ 0.01 at day 21 by paired t-test), and volumes of untreated tumors on day 15 (I, *right*, **P ≤ 0.01; **** P ≤ 0.0001 by one-way ANOVA). **(J)** Kaplan-Meier survival curves for mice with two B16.F10-OVA tumors. Tumors were measured by a caliper every other day from day 7 through 30 after tumor implantation (n = 7-10 mice per group). All data are presented as mean ± SD.

We next evaluated the effect of SLR-LNPs on tumor growth using the B16.F10 melanoma model, employing an intratumoral administration (i.t.) route that is used clinically in melanoma patients receiving oncolytic virus therapy (e.g., T-VEC) (**Figure 2C**).^30^ We found that SLR-LNPs inhibited tumor growth, resulting in an increase in survival time, whereas empty LNPs and cSLR-LNPs had no impact on tumor growth inhibition relative to vehicle (PBS) treated mice (**Figure 2D-F**). We also tested therapeutic efficacy in an ovalbumin-expressing B16.F10 (B16-OVA) model in which two tumors were established on opposite flanks and only one of the tumors (treated) was injected with SLR-LNPs (**Figure 2G**). Consistent with induction of an abscopal effect, we found that i.t. administration of SLR-LNP inhibited growth of the treated tumor (**Figure 2H**), but also reduced growth of the distal (untreated tumor) (**Figure 2I**). As was observed in the single tumor study, empty LNPs and cSLR-LNP had no effect on tumor growth inhibition in this dual-tumor model, indicating that the antitumor response is RIG-I dependent. We also evaluated SLR-LNPs in combination with αPD-1 and αCTLA4 ICIs, which are approved for the treatment of metastatic melanoma. As expected in the immunologically cold B16-OVA model, αPD-1 + αCTLA4 had no effect on tumor growth but enhanced the efficacy of SLR-LNPs in inhibiting both primary and distal tumor growth and increasing overall survival (**Figure 2G-J**). These studies demonstrate that intratumoral administration of SLR-LNPs can inhibit both treated and distal tumor growth and increase response to approved ICIs approved in melanoma. While other materials (e.g., jetPEI) have been employed for local delivery of RIG-I agonists,^16, 24^ it is notable that LNPs have now been locally administered via intramuscular injection to millions of people receiving COVID-19 mRNA vaccines, which may accelerate the translation of SLR-LNPs for intralesional therapy.

### Systemic administration of SLR-LNPs inhibits tumor growth

While SLR-LNPs hold promise as an intralesional therapy, intratumoral administration may not be possible or practical for many patients and/or cancer types.^31^ Therefore, we next focused our investigations on the larger challenge of achieving safe and effective systemic administration of RIG-I agonists for cancer immunotherapy. Our group has recently identified RIG-I activation as a potentially promising target for enhancing immunotherapy responses in triple negative breast cancer (TNBC).^28^ Therefore, we evaluated the efficacy of systemically administrated SLR-LNPs in an orthotopic EO771 breast cancer model. We first administered SLR-LNPs or control formulations intravenously at a dose of 10 μg SLR (∼0.5 mg/kg) three times, spaced three days apart, and monitored tumor volume (**Figure 3A**). SLR-LNPs significantly inhibited tumor growth and increased survival time, whereas empty LNPs and cSLR-LNPs had no effect (**Figure 3B,C**), further demonstrating the importance of RIG-I activation in mediating a therapeutic benefit. We further tested the efficacy of intravenously administered SLR-LNPs in the B16.F10 model, again observing inhibition of tumor growth and extended survival time (**Figure 3D-F**).

**Figure 3.**
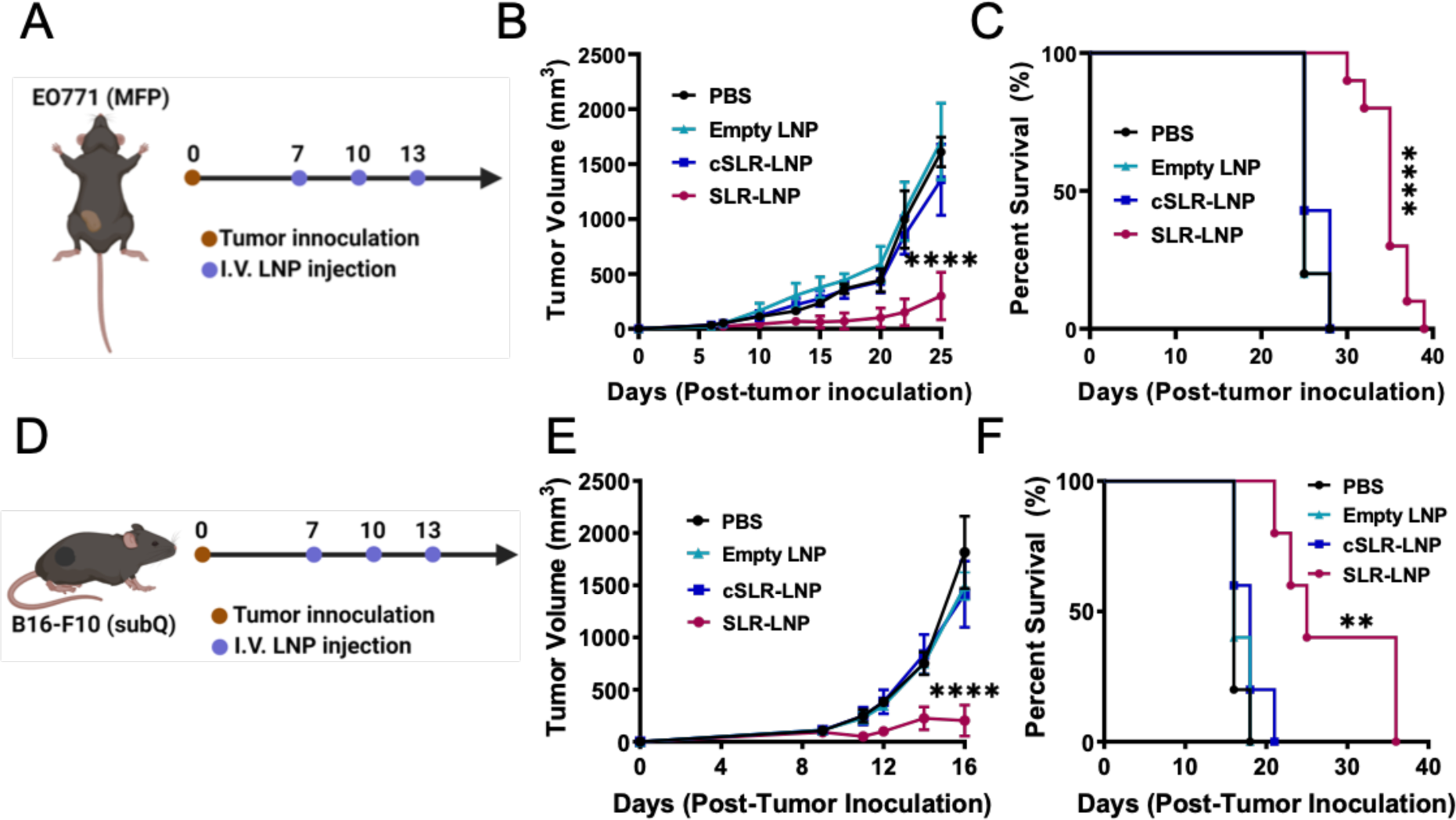
Systemic administration of SLR-LNPs inhibits tumor growth. **(A)** Schematic illustration of intravenous LNP treatment in mice bearing EO771 tumors in the mammary fat pad (MFP). **(B)** Tumor growth curves of EO771 tumor-bearing mice after the indicated treatments. (n= 8-10 per group, ****P ≤ 0.0001 by one-way ANOVA compared to all other groups). **(C)** Kaplan-Meier survival curves for EO771 MFP tumor-bearing mice after the indicated treatments. **(D)** Schematic illustration of intravenous LNP treatment in B16-F10 tumor-bearing mice. **(E)** Tumor growth curves of B16-F10 tumor-bearing mice after the indicated treatments. (n=5 per group, ****P ≤ 0.0001 by one-way ANOVA compared to all other groups). **(F)** Kaplan-Meier survival curves for B16-F10 tumor-bearing mice after the indicated treatments. All data are presented as mean ± SD.

Importantly, we found that this therapeutic regimen was well tolerated, with mice exhibiting only a mild (∼5%) and transient weight loss in the immediate post-treatment period (**Figure S1B**). Consistent with administration of other innate immune agonists, including those that have advanced into the clinic,^18, 32, 33^ elevated serum cytokine levels were observed six hours following administration but were insignificant from background by 24 h (**Figure S1C**). Additionally, no changes in levels of serum BUN, ALT, glucose, or AST were observed (**Figure S1D**), indicating that the treatment did not induce significant liver or kidney damage. Red blood cell (RBC) count and hemoglobin (HGB) levels were slightly reduced for all nanoparticle formulations, but no effect on mean corpuscular hemoglobin (MCH) or MCH concentration (MCHC) were noted. Complete blood count (CBC) revealed no differences relative to vehicle control, with the exception of neutrophils, which were elevated in response to SLR-LNP treatment. Major organs (liver, spleen, kidney, lung, brain, heart, pancreas, and bone marrow (sternum)) were also isolated 24 h following treatment, routinely fixed in 10% neutral buffered formalin, embedded, sectioned, and stained for blinded evaluation by a board-certified veterinary pathologist. No histopathologic abnormalities were observed in the kidney, lung, brain, heart, or pancreas. Histological evidence of a slight increase in extramedullary hematopoiesis in the liver and spleen was observed, and an increased ratio of myeloid to erythroid precursor cells in the bone marrow was noted, both of which were likely secondary to elevated pro-inflammatory cytokine levels and not clinically significant.

### Systemic administration of SLR-LNPs reprograms the TME to enhance T cell infiltration

Having established a safe and effective regimen for systemic administration of SLR-LNPs, we next evaluated effects on the tumor microenvironment in the orthotopic EO771 breast cancer model. Tumor tissue was harvested 24 h following the three-dose regimen and processed for analysis by qRT-PCR, western blot, and flow cytometry (**Figure 4A**). Consistent with RIG-I activation, we observed increased levels of pIRF3 in the TME (**Figure 4B**) and expression of ISGs and proinflammatory cytokines (**Figure 4C**) in mice treated with SLR-LNPs, but not empty LNPs or cSLR-LNP formulations. We also observed a significant increase in the number of tumor infiltrating CD4^+^ and CD8^+^ T cells in response SLR-LNP treatment, but no significant differences in the number of NK cells, dendritic cells, macrophages, or MDSCs (**Figure 4D**). Based on these data, we antibody depleted CD4^+^ and CD8^+^ T cells to elucidate their relative contributions to antitumor efficacy in the EO771 model (**Figure 4E-G**). Depletion of either T cell population abrogated therapeutic efficacy, with CD8^+^ T cell depletion having a slightly larger impact on antitumor efficacy (**Figure 4F-G**). Collectively, these studies demonstrate the ability of SLR-LNPs to promote the infiltration of T cells with antitumor function into the TME.

**Figure 4.**
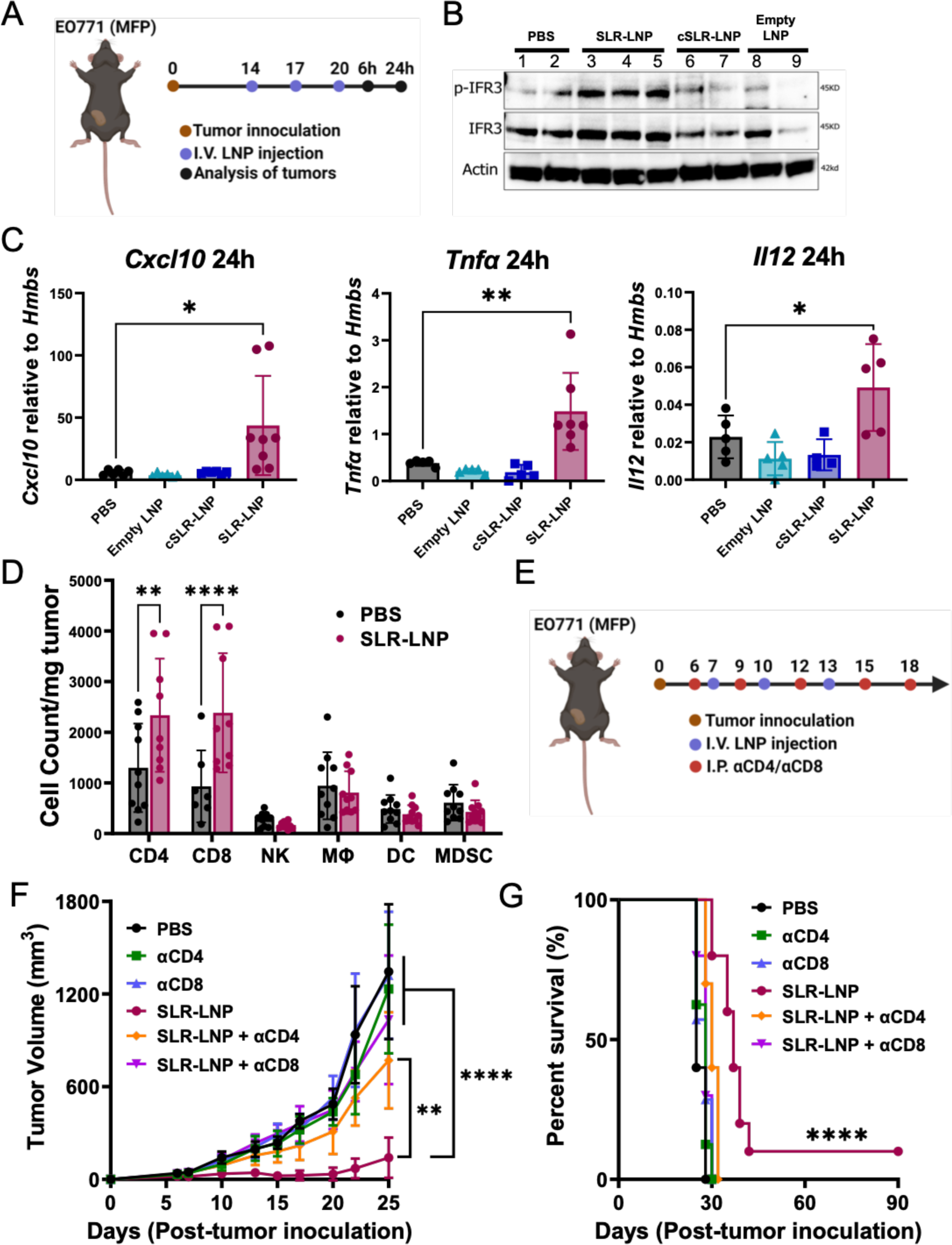
Systemic administration of SLR-LNPs activates RIG-I in the tumor microenvironment to enhance infiltration of CD8^+^ and CD4^+^ T cells with antitumor function. **(A)** Schematic diagram of LNP treatment in the orthotopic EO771 breast cancer model. **(B)** The effects of SLR-LNP on RIG-I signaling were determined using Western blot analysis. **(C)** RT-PCR analysis of pro-inflammatory cytokines gene expression in the tumor. **P≤0.01, ***P ≤ 0.001 by one-way ANOVA. **(D)** Flow cytometric analysis of immune cells in the TME including CD8^+^ and CD4^+^ T cells, macrophages (CD11b^+^F4/80^+^), dendritic cells (MHC-II^+^CD11c^+^), NK cells (NK1.1^+^) and MDSCs (CD11b^+^Gr^+^). **P ≤ 0.01, ****P ≤ 0.0001 by 2-way ANOVA. **(E)** Schematic diagram of experimental design in EO771 MFP tumors treated with anti-CD4 (αCD4) or anti-CD8 (αCD8) depleting antibodies, and SLR-LNP (n = 8-10 mice per group). **(F)** Comparison of tumor growth curves (**P ≤ 0.01, ****P ≤ 0.0001 by one-way ANOVA of tumor volumes at day 25) and **(G)** Kaplan-Meier overall survival curves after T cell depletion and SLR-LNP treatment in EO771 MFP tumor-bearing mice. All data are presented as mean ± SD.

### SLR-LNPs enhance response to immune checkpoint blockade

Based on the capacity of SLR-LNPs to promote T cell infiltration into the breast TME, we next evaluated SLR-LNPs in combination with αPD-1 ICI, which is approved for a subset of TNBC patients, who experience a response rate of only ∼20%.^34^ Mice with orthotopic EO771 tumors were treated with SLR-LNPs alone or in combination with anti-PD-1 ICI and tumor volume was monitored (**Figure 5A**). SLR-LNPs inhibited tumor growth to a greater degree than αPD-1, which exerted only minimal efficacy as monotherapy (**Figure 5B**), and the combination of SLR-LNPs and αPD-1 further inhibited tumor growth and extended survival, resulting in a 25% (2/8) complete response rate (**Figure 5C**). We also tested SLR-LNPs in the context of aggressive metastatic melanoma, a setting where systemic administration of RIG-I agonists may be necessary. As a model of lung metastasis, luciferase-expressing B16.F10 cells were injected intravenously to colonize the lung, and mice were subsequently treated with SLR-LNP alone and in combination with αCTLA-4/αPD-1 ICI (**Figure 5D**). Mice were euthanized 20 days post-tumor inoculation and lung metastatic burden was evaluated via luminescence and lung mass measurements. Consistent with our other findings, SLR-LNPs, but not cSLR-LNPs, dramatically inhibited tumor formation in the lung, an effect that was further, though not significantly, enhanced with the addition of αCTLA-4/αPD-1 ICI, which had no effect as a monotherapy in this model (**Figure 5E-H**). Hence, systemic administration of SLR-LNPs can inhibit tumor growth and metastasis as well as increase response to approved ICIs in multiple poorly immunogenic tumor models.

**Figure 5.**
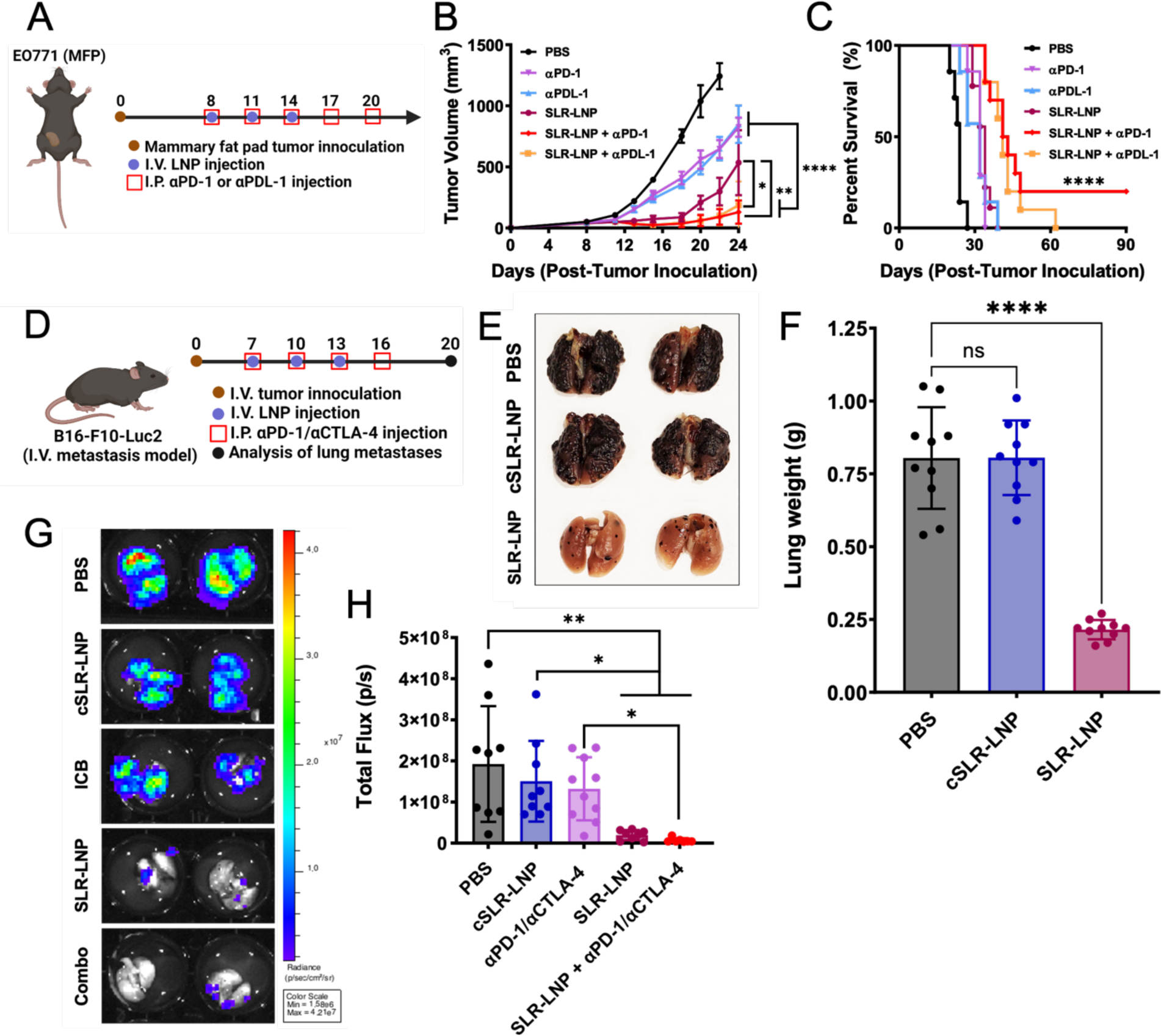
SLR-LNPs enhance response to immune checkpoint inhibitors. **(A)** Schematic diagram of LNP and anti-PD-1antibody (αPD-1) treatment in the orthotopic EO771 model (n=8-10). **(B)** Comparison of tumor growth curves (*P ≤ 0.05, ***P ≤ 0.001, ****P ≤ 0.0001 by one-way ANOVA of tumor volumes at day 24, ####P≤ 0.0001 compared to all other groups on day 22 by one-way ANOVA). **(C)** Kaplan-Meier overall survival curves after SLR-LNP and αPD-1 treatment in EO771 MFP tumor-bearing mice. **(D)** Schematic diagram showing the schedule for B16F10-Luc1 tumor challenge and treatment (n=10). **(E)** Representative lung images from PBS, cSLR-LNP, and SLR-LNP treatment groups. **(F)** Quantification of lung weight (****P ≤ 0.0001 by one-way ANOVA). **(G)** Bioluminescence images measured by IVIS of lungs. **(H)** Quantification of lung metastases by FLI depicted in (G) (*P ≤ 0.05, **P ≤ 0.01 by one-way ANOVA.) All data are presented as mean ± SD.

## Discussion

Identifying agents that remodel the tumor microenvironment from “cold” (i.e., lacking T cell infiltration) to “hot” (i.e., T cell inflamed) has rapidly emerged as a promising approach for reversing resistance to ICIs.^5, 6, 35, 36^ RIG-I is promising target for increasing tumor immunogenicity and improving response to immunotherapy, but major pharmacological barriers limit the activity and efficacy of 3pRNA as a nucleic acid therapeutic. Here, we address this challenge using a facile and translationally-ready strategy that leverages advanced LNP technology and a molecularly engineered SLR to fabricate an immunotherapeutic nanomedicine for potent activation of RIG-I signaling. We found that SLR-LNPs, administered either intratumorally or intravenously, activated RIG-I in the tumor microenvironment, resulting in enhanced effector T cell infiltration that inhibited tumor growth and enhanced response to ICIs in multiple immunologically “cold” tumor models. This represents a new application of LNPs and establishes a foundation for further optimization and preclinical development of SLR-LNPs for cancer immunotherapy.

Despite the promise of 3pRNA as an immunopotentiator, there has been relatively little investigation into the design and testing of carriers to enhance its efficacy. Several groups have employed PLGA-based micro- and nanoparticle formulations for 3pRNA delivery, primarily for vaccine applications.^37, 38^ Huang and co-workers described 3pRNA-loaded lipid calcium phosphate nanoparticles and demonstrated that systemic administration could inhibit tumor growth in models of pancreatic cancer.^39^ Our has described polymeric carriers for 3pRNA delivery by exploiting the combinatorial diversity enabled through the synthesis of polymer and SLR structural libraries.^17, 40^ However, recent clinical trials,^25^ and most preclinical studies to date^11, 13, 23, 24^, have employed the cationic polymeric transfection reagent JetPEI^TM^, which electrostatically condenses nucleic acids and facilities their release from the endolysosome.^41^ While PEI-based approaches remain promising, and merit continued development, their clinical translation has been hindered by toxicity concerns, a proclivity for accumulation in the lungs, and a relatively low efficiency for cytosolic delivery via the still debated “proton sponge” mechanism.^42, 43^ By contrast, LNPs have rapidly emerged as one of the most versatile platforms for delivery of a diverse array of nucleic acids, and are essential to the efficacy of several recently approved nucleic acid therapeutics.^26^ Additionally, LNPs are approved for administration both locally (e.g., as mRNA vaccines) and intravenously (e.g., as siRNA therapeutics), providing a versatile drug carrier for both intralesional therapy and systemic therapy for treatment of metastatic disease.

Surprisingly, there has been virtually no investigation into the use of LNPs for delivery of 3pRNA, which faces delivery barriers common to other classes of oligonucleotide therapeutics but is distinguished by its unique immunopharmacological mechanisms of action. Hence, we sought to fill this knowledge and innovation gap by leveraging LNP technology to design and test a nanoscale platform for RIG-I activation. Our selection of the MC3 ionizable lipid was primarily motivated by translational considerations as it is already approved for clinical use, and, therefore, represented a logical initial choice for the design RIG-I activating lipid nanoparticles. However, there is also now a vast tool box of ionizable lipids available for RNA delivery that vary in head group and lipid tail structure that can be leveraged to optimize delivery of a specific nucleic acid cargo.^26^ Further, LNP formulations can be assembled with different types and/or compositions of helper and PEGylated lipids using different fabrication approaches, which can be harnessed to modulate pharmacokinetics and/or to confer tissue-or cell-specific tropism that can be tuned for specific disease applications.^44^ Hence, there is a virtually limitless parameter space for the design of LNPs for the delivery of 3pRNA that can now be explored by building upon our development and evaluation of a first-generation SLR-LNP platform for immunotherapy.

The design of drug carriers for 3pRNA therapeutics will ultimately be driven by an understanding of pharmacological mechanisms of efficacy and toxicity. Such knowledge remains limited for this unique class of oligonucleotide therapeutic due, in part, to a dearth of technologies that have been developed and/or tested for 3pRNA administration. In this regard, our investigations provide a preclinical benchmark for evaluating systemically administered RIG-I agonists and their carriers. An important distinction between 3pRNA and more conventional classes of nucleic acid therapeutics for cancer (e.g., siRNA) is that 3pRNA can exert robust therapeutic effects via both local and systemic reprogramming of immune cell populations that can initiate and propagate antitumor immunity. This has the potential to obviate the need to deliver high drug doses to the vast majority of cancer cells at all tumor sites in the body, which is an established limitation of even the most promising nanomedicines.^45^ While our study demonstrates that SLR-LNPs can enhance T cell infiltration into tumors and that the response is T cell-mediated, further investigation is necessary to identify the primary cellular responders to SLR-LNPs and to examine the immunopharmacological behavior of SLR-LNPs in the tumor and other tissues (e.g., liver, spleen).

While intravenous administration of SLR-LNPs results in RIG-I activation in the TME, it also results in a transient elevation of serum cytokines due to on-target, off-tumor effects. While systemic immunopotentiation that mobilizes an antitumor response may be advantageous, and perhaps even necessary, for the treatment of advanced metastatic disease, the potential of systemically administrated PRR agonists to induce inflammatory toxicities must be considered.^18^ Therefore, we performed a robust preclinical analysis of toxicity following a therapeutic three-dose regimen and found SLR-LNPs to be well-tolerated, with mice exhibiting only mild and transient weight loss without evidence of organ pathology or abnormal blood chemistry test results. It is notable that other promising innate immune agonists similarly induce a transient elevation of serum cytokines and weight loss in mice,^46, 47^ including nanomedicines and antibody-drug conjugates that have advanced into patients, which experienced only transient flu-like symptoms or other adverse events that were readily manageable.^33, 48^ Nonetheless, an important future direction will be to engineer SLR-LNPs to further enrich RIG-I activation in the TME, while minimizing systemic inflammatory responses. Towards this end, our group has described the design of 3pRNA pro-drugs that employ bulky covalently-linked macromolecules (e.g, PEG) to block RIG-I recognition of 3pRNA until they are removed under a specific environmental stimulus (e.g., redox, enzymes).^49^ Likewise, there is deep nanomedicine toolbox available for improving cargo delivery to tumor sites, including integration of molecular targeting moieties or “sheddable” coronas, that could be harnessed to expand the therapeutic window of systemically administered RIG-I agonists. While systemically administered SLR-LNPs demonstrated efficacy as a monotherapy, their ability to remodel the TME to increase CD8^+^ and CD4^+^ T cell infiltration also offers exciting opportunities for developing novel combination immune-regimens to further enhance therapeutic responses. Here, we focused our investigations on combining SLR-LNPs with approved ICIs based on the recent Phase I clinical trials that explored intratumoral injection of 3pRNA in combination with anti-PD-1 (pembrolizumab).^25^ But there is also a strong immunological rationale for combining RIG-I agonists with other approved and experimental therapeutics, including chemotherapy and other immunomodulators. Furthermore, our finding that SLR-LNPs increase the tumor infiltration of endogenous T cells opens the possibility of using i.v. administered SLR-LNPs to enhance responses to other T cell-based immunotherapeutic modalities – including cancer vaccines, adoptive T cell transfer, or CAR T cell therapy – where poor tumor infiltration is a major barrier to efficacy for solid tumors.^37, 50^ SLR-LNPs pave the way to pursue these opportunities.

## Conclusion

In conclusion, we have described the fabrication, characterization, and preclinical evaluation of a nanoparticle-based immunotherapy that enhances antitumor immunity via activation of the RIG-I pathway. Our design of a nanoparticle RIG-I agonist was inspired by currently approved lipid nanoparticle formulations for other classes of RNA therapeutics: we leveraged the ionizable lipid DLin-MC3-DMA to package and enhance the intracellular delivery of selective and well-defined 5’-triphosphate stem loop RNA (SLR) RIG-I ligands. We demonstrated that this strategy resulted in potent activation RIG-I signaling *in vitro* and *in vivo*, and that SLR-LNPs could be safely administered via both intratumoral and intravenous routes to promote RIG-I activation in the TME, resulting in expression of type-I interferons, proinflammatory cytokines, and chemokines that enhanced the infiltration of CD8^+^ and CD4^+^ T cells with antitumor function. Consequently, SLR-LNPs inhibited tumor growth in a RIG-I-dependent manner in multiple poorly immunogenic solid tumor models and increased therapeutic responses to anti-PD-1 and anti-CTLA-4 immune checkpoint inhibitors. Collectively, these studies establish lipid nanoparticle-based delivery of RIG-I agonists as a translationally promising strategy for increasing tumor immunogenicity and enhancing responses to ICs and other immunotherapies.

## Materials and Methods

### Synthesis of DLin-MC3-DMA lipid

DLin-MC3-DMA (MC3) was prepared following the method described in WO2010144740 (Example 5, page 140). Detailed synthesis methods are available in Supporting Information and characterization by 1H NMR, UPLC-ELSD, and mass spectrometry is provided in **Figure S2**.

### Formulation of SLR-LNPs

SLR20 was synthesized and purified as described previously.^17, 24^ LNP formulations of SLR20 were prepared as previously described for formulation of siRNA-loaded LNPs with minor modifications.^51^ Briefly, DLin-MC3-DMA, 1,2-distearoyl-sn-glycero-3-phosphocholine (DSPC) (Avanti Polar Lipids), cholesterol (Avanti Polar Lipids), and PEG_2kDa_-lipid (PEG-DMG) (Avanti Polar Lipids) were solubilized in ethanol at a molar ratio of 57.5:7.5:31.5:3.5 and heated to 65°C prior to dropwise addition into citrate buffer (0.1 M, pH 3, 25°C) under constant mixing to a final volume ratio of 1:3 ethanol to citrate buffer. For SLR-containing formulations, SLRs were dissolved in citrate buffer prior to lipid addition at a concentration that resulted in a final SLR weight fraction (w/w) of 0.06 SLR/Dlin-MC3-DMA; for *in vivo* studies, an SLR weight fraction of 0.1 was used. Homogenous mixing was allowed to occur for at least 1 h at room temperature to ensure nanoparticle formation. The ethanol and citrate were removed via buffer exchange with PBS (155 mM NaCl, 3 mM Na_2_HPO_4_, 1 mM KH_2_PO_4_, pH 7.4) by dialysis using an Amicon^®^ Ultra-15 Centrifugal Filter Unit with 100 kDa molecular weight cutoff regenerated cellulose membrane (Millipore) or via tangential flow filtration (Repligen; KrosFlo Research I Peristaltic Pump with MicroKros Hollow Fiber Filter) for larger batches used for mouse studies. Particle size and zeta potential were determined using a Malvern Zetasizer Nano ZS instrument at room temperature. The measurement was repeated three times independently for each sample. The amount of encapsulated nucleic acid was determined using the Quant-it™ RiboGreen RNA Assay Kit (Invitrogen). Briefly, LNPs were disrupted in 2% Triton X-100 in TE buffer, RiboGreen solution was added to these samples, and fluorescence was measured using a plate reader (Synergy H1 Multi-Mode Microplate Reader; Biotek). The RNA concentration was then determined by comparing the fluorescence of the LNP samples to SLR20 or SLROH standard curves.

### Cell culture

B16-F10 cells were obtained from American Type Culture Collection (ATCC) (Manassas, Virginia) and RAW-Dual ISG, THP1-Dual ISG, A549-Dual ISG, RAW-Dual RIG-I KO ISG were purchased from InvivoGen. EO771 cells were gifted from Justin Balko (Vanderbilt University Medical Center), luciferase-expressing B16-F10 cells (B16-LUC) were provided by Ann Richmond (Vanderbilt University Medical Center), and ovalbumin expressing B16-F10 (B16-OVA) cells were gifted from Amanda Lund (New York University School of Medicine). B16-F10 cells expressing an interferon inducible luciferase reporter were used as described previously.^46^ All cell lines were cultured according to manufacturer’s specifications. BMDCs were isolated from 6-8-week-old C57BL/6 mice and cultured as previously described.^52^ Briefly, bone barrow was flushed from the femurs and tibias of mice using complete BMDC culture medium (RPMI 640 medium supplemented with 10% heat-inactivated FBS, 100 U/mL penicillin, 100 μg/mL streptomycin, 2 mM l-glutamine, 10 mM HEPES, 1 mM sodium pyruvate, 1× non-essential amino acids, and 50 μM β-mercaptoethanol), and the marrow was passed through a 70 μM cell strainer. Cells were centrifuged at 1500 rpm for 5 minutes, resuspended in ACK lysis buffer (ThermoFisher), and washed with cold PBS. Then, cells were seeded in 100 × 15 mm Petri dishes (Corning Inc.) in complete medium supplemented with 20 ng/mL granulocyte-macrophage colony-stimulating factor (GM-CSF). Cells were maintained in a 37°C incubator supplemented with 5% CO_2_, and culture medium containing GM-CSF was replaced on days 3, 5, and 7. On day 8, cells were confirmed to be > 80% BMDCs (CD11c^+^) by flow cytometry.

### Evaluation of immunostimulatory activity in ISG reporter cells

RAW-Dual ISG, THP1-Dual ISG, A549-Dual ISG, and RAW-Dual RIG-I KO ISG Reporter cells were seeded at 5×10^5^ cells/well in 100 µL media in 96-well plates (Greiner Bio-One). When adherent cells became ∼80% confluent or suspension cells reached a density of 1.5×10^6^ cells/mL, SLR-LNPs or controls were added to wells at 2x concentration in 100 µL media. Supernatant was collected 24h after treatment and Quanti-Luc^TM^ (Invivogen) assay used to determine the amount of secreted luciferase following manufacturer’s instructions. Luminescence was quantified using a plate reader (Synergy H1 Multi-Mode Microplate Reader; Biotek) using white, opaque-bottom 96-well plates (Greiner Bio-One). The signal for each sample concentration was determined using 3 biological replicates, each with 3 technical replicates. All reporter cell measurements were normalized by subtracting the average value of a PBS-treated negative control group. The EC_50_ values were calculated for each of the dose responses using curve fitting analysis in the GraphPad Prism software.

### Gene expression in BMDCs and cancer cell lines

Relative gene expression of *Ifnβ1, Tnfα, Cxcl10* and/or *IL12* in BMDCs, B16.F10 melanoma cells, and EO771 breast cancer cells was quantified by qPCR following treatment with SLR-LNP or controls. In brief, 1 × 10^6^ BMDCs/well, 500,000 B16.F10 cells/well, or 500,000 EO771 cells/well were seeded in 12-well plate and treated with PBS, empty LNP cSLR-LNP, or SLR-LNP for 24 hrs. Total RNA was isolated using a RNeasy Mini kit (Qiagen, Germantown, MD). Total RNA (1 μg) was reverse transcribed by an iScript cDNA synthesis kit (Bio-Rad) and qPCR was performed using a TaqMan Mastermix kit (Thermo Fisher Scientific) as per manufacturer’s instructions.

### Evaluation of BMDC activation

BMDC activation was evaluated by flow cytometric analysis of surface CD80, CD86, and MHC-II expression. Briefly, 1 × 10^6^ BMDCs/well were seeded in a 12-well plate and treated with PBS, empty LNP, cSLR-LNP, or SLR-LNP for 24 hrs. The cells were collected and washed with 3% BSA in PBS and then stained with FITC-anti-CD11c (1:100), APC/Cy7-anti-MHC class II (1:100), PE-anti-CD86 (1:100), APC-anti-CD80 (1:100) (Biolegend) antibodies. Dead cells were excluded from analysis using DAPI (live/dead) stain (1:20,000). Cells were analyzed using a CellStream flow cytometer.

### Animal ethics statement

All studies using animals were completed under an Animal Care Protocol approved by the Vanderbilt University Institutional Animal Care and Use Committee (IACUC). Animal health assessments were completed using standard operating procedures approved by Vanderbilt University IACUC.

### Subcutaneous single B16-F10 tumor model

B16-F10 (3×10^5^) cells were subcutaneously injected into the right flank region of 6-7-week-old female C57BL/6 mice (The Jackson Laboratory, Bar Harbor, ME). Established B16-F10 (40-60 mm^3^) tumors were treated intratumorally with vehicle (PBS), empty LNP, cSLR-LNP (10 µg), SLR-LNP (10 µg) in 50 μL. For evaluation of gene expression via qPCR, mice were treated once intratumorally and mice were euthanized 24 h post-injection. For evaluation of therapeutic efficacy, mice were administered SLR-LNPs or controls intratumorally every 3 days for 3 total injections. Tumor volume was measured 3x weekly via caliper measurements, and volumes were calculated using (V_tumor_ = L × W^2^ × 0.5, in which V_tumor_ is tumor volume, L is tumor length, and W is tumor width). Mice were euthanized by carbon dioxide asphyxiation when tumor volume reached >1500 mm^3^.

### Orthotopic EO771 breast tumor model

EO771 (2.5×10^5^) cells were injected into the left inguinal mammary fat pad of 6-7-week-old female C57BL/6 mice (The Jackson Laboratory, Bar Harbor, ME). Mice were randomized into treatment groups and mice were intravenously administered vehicle (PBS), empty LNP, cSLR-LNP (10 µg), or SLR-LNP (10 µg) in 100 µL PBS 3 times spaced 3 days apart. In studies evaluating effects on the tumor microenvironment by qRT PCR, western blot analysis, or flow cytometry, mice were euthanized 24 h following the last treatment. In studies investigating combination effects with immune checkpoint inhibitors (αPD-1) or T cell depletion antibodies (αCD4 or αCD8), mice received intraperitoneal injections of 100 µg antibody in dilution buffer every 3 days for 5 total injections. For T cell depletion studies, antibody treatment began 24 h before treatment with SLR-LNPs. Tumor volume was measured 3x weekly via caliper measurements, and volumes were calculated using (V_tumor_ = L × W^2^ × 0.5, in which V_tumor_ is tumor volume, L is tumor length, and W is tumor width). Mice were euthanized by carbon dioxide asphyxiation when tumor volume reached >1500 mm^3^.

### *In vivo* imaging of interferon response

B16.F10 melanoma cells were transduced to express luciferase in an ISRE-dependent manner via the Cignal Lenti Reporter construct (Qiagen) as described previously.^46^ 6-8 week-old C57BL/6 mice (The Jackson Laboratory) were anesthetized with isoflurane and their right dorsal flanks were shaved. Mice were inoculated with 1 × 10^6^ B16.F10 interferon reporter cells in 100 µL of PBS. When tumors were ∼100 mm^3^, the mice were administered a single 50 µL intratumoral injection of either PBS or SLR-LNP at a dose corresponding to 10 µg SLR20. At each timepoint (0 and 6 h), mice were administered a dorsal subcutaneous 150 µL injection of 30 mg/mL D-luciferin (Thermo Fisher Scientific) reconstituted in PBS and luminescence images was captured 15 minutes thereafter. Relative IFN production for each tumor was calculated at 6h as a fold change relative to the respective t=0h value for each mouse.

### Subcutaneous B16-OVA two tumor model

Ovalbumin-expressing B16-F10 melanoma (B16-OVA) cells (2.5×10^5^) cells were subcutaneously injected into the right flank region of 6-7-week-old female C57BL/6 mice (The Jackson Laboratory, Bar Harbor, ME). A second subcutaneous injection containing (1.5×10^5^) cells was performed on the left flank region 2 days after the initial tumor inoculation. Established right flank B16-OVA (40-60 mm^3^) tumors were treated intratumorally with vehicle (PBS), empty LNP, cSLR-LNP or SLR-LNP (10 µg) every 3 days for 3 total injections. Mice in groups receiving αPD-1 and αCTLA-4 (100 µg, every 3 days for 5 injections, BioXcell, West Lebanon NH) were treated intraperitoneally. Tumor volume was measured 3x weekly via caliper measurements, and volumes were calculated using (V_tumor_ = L × W^2^ × 0.5, in which V_tumor_ is tumor volume, L is tumor length, and W is tumor width). Mice were euthanized by carbon dioxide asphyxiation when tumor either reached >1500 mm^3^.

### Lung metastatic B16-F10 tumor model

6–8 week-old C57BL/6 mice (The Jackson Laboratory) were administered a single intravenous injection of 0.5 × 10^6^ luciferase-expressing B16.F10 cells (B16-LUC) suspended in PBS. On day 3 post tumor inoculation, mice were treated intravenously with PBS, cSLR-LNP (10 µg), or SLR-LNP (10 µg) every 3 days for 3 doses total. Mice in groups receiving αPD-1 and αCTLA-4 (100 µg, every 3 days for 4 injections, BioXcell, West Lebanon NH) were treated intraperitoneally. Twenty days post tumor inoculation, mice were euthanized, and lungs were excised. Lungs were weighed and imaged. Lungs were then placed in black 12-well plates (Cellvis) and incubated in 1 mg/mL Pierce™ D-Luciferin, Monopotassium Salt (88293; Thermo Fisher Scientific) reconstituted in PBS, and luminescence images were captured 5 minutes thereafter on the IVIS Lumina III (PerkinElmer). The luminescence was quantified as total radiant flux (p/s) for each set of lungs.

### Quantitative real-time PCR (qPCR) of tumor tissue

C57BL/6 mice bearing either subcutaneous B16-F10 or orthotopic EO771 breast tumors were treated with SLR-LNPs or controls as described above and euthanized 6 or 24 h later. Tumors were collected and snap frozen in liquid nitrogen until analysis. Tumors were homogenized using a TissueLyser II (Qiagen) and total RNA was isolated by RNeasy mini kit (Qiagen) and reverse transcribed by iScript cDNA synthesis kit (Bio-Rad). qPCR was performed using a TaqMan Mastermix Kit (Thermo Fisher Scientific) as per manufacturer’s instructions. *Hmbs* was used as a housekeeping gene. TaqMan gene expression primers were purchased from ThermoFisher Scientific (Waltham, Massachusetts): mouse *Tnfa* (Mm00443258_m1); mouse *Cxcl10* (Mm00445235_m1); mouse *Il12b* (Mm00434174_m1) and mouse *Hmbs* (Mm01143545_m1).

### Western blot analysis of EO771 tumors

Female C57BL/6 mice with 100-200 mm^3^ EO771 tumors in the mammary fat pad were treated as described above with SLR-LNP or controls. Mice were euthanized at 24 h following the last injection and tissues were snap frozen until analysis. EO777 tumors were homogenized using a TissueLyser II (Qiagen) in RIPA lysis buffer (Santa Cruz). The protein concentration was determined using a BCA assay (Thermo Scientific, Waltham, Massachusetts), samples were run on a SDS-PAGE and transferred onto a nitrocellulose membrane using a semi-dry transfer protocol (Bio-Rad laboratories, Hercules, California). Membranes were washed and incubated with primary antibody (p-IRF3, IRF3, RIG-I, and β-actin) at 4°C overnight, followed by blotting with HRP-conjugated secondary antibodies (Promega). The protein bands were obtained with the ChemiDoc XRS+system (Bio-Rad) using an immobile western Chemiluminescent HRP Substrate Kit (Millipore Sigma, Billerica, Massachusetts). Protein from blots was quantified using ImageJ, and β-actin was used as a loading control for normalization of samples.

### Flow cytometric analysis of EO771 tumors

Female C57BL/6 mice with 100-200 mm^3^ EO771 tumors in the mammary fat pad were treated with SLR-LNPs (10 µg, intravenously) or PBS every 3 days for three total injections. Mice were euthanized 24 h after final treatment, tumors were harvested, weighed, and placed on ice prior to dissociation using an OctoMACS separator (Miltenyi) and digestion in RPMI 1640 containing 125 µg/mL deoxyribonuclease I and 500 µg/mL collagenase III for 30 mins at 37°C. The cell suspension was strained through a cell strainer (40 µm), red blood cells were lysed using ACK lysis buffer (Gibco). Cells were then resuspended in flow buffer (5% BSA + 0.1% dasatinib in PBS), counted and stained with the following flow panels (antibodies obtained from Biolegend). Panel I for T cells: PE/Cy7-αCD45 (30-F11), APC-αCD3 (17A2), APC/Cy7-αCD4 (RM4–5), PE-αCD8α (53-6.7) and DAPI. Panel II for myeloid cells and NK cells: APC-αCD45 (30/F11), PerCP/Cy5.5-αCD11b (M1/70), PE/Cy7–αF4/80 (BM8), Alex Flour 488-αCD11c (N418), APC/Cy7-αMHC-II (M5/114.15.2), PE-αNK1.1 (PK136), BV605-Gr-1 (1A8), and DAPI (Millipore Sigma, Billerica, Massachusetts). Cells were then washed twice, suspended in flow buffer containing AccuCheck counting beads and analyzed on a BD LSR II flow cytometer. All flow cytometry data were analyzed using FlowJo software (version 10; Tree Star; https://www.flowjo.com/solutions/flowjo). Representative flow cytometry plots and gating schemes are shown in **Figure S3**.

### Statistical Analysis

The data were plotted using Prism 8 (Graphpad) software as the mean ± SD unless otherwise stated in the figure legend. Data were analyzed via Student’s t-test or a one-way ANOVA followed by Tukey’s adjustment for multiple comparisons. A log-rank test was used to compare Kaplan-Meier survival data. P-values <0.05 were considered statistically significant in all studies.

## Acknowledgements

The authors would like to thank Dr. Justin Balko for donating EO771 cells, Dr. Ann Richmond for providing B16-LUC cells, Dr. Amanda Lund for the gift of B16-OVA cells, and Dr. Craig Duvall for use of IVIS Imaging System. We thank the core facilities of the Vanderbilt Institute of Nanoscale Sciences and Engineering (VINSE) and the VUMC Flow Cytometry Shared Resource, supported by the Vanderbilt Ingram Cancer Center (P30 CA68485) and the Vanderbilt Digestive Disease Research Center (DK058404). This research was supported by grants from the CDMRP Breast Cancer Research Program (W81XWH-20-0624 to JTW), CDMRP Kidney Cancer Research Program (W81XWH-22-1-0661 to JTW), a Vanderbilt Ingram Cancer Center (VICC) Ambassador Discovery Grant (JTW), and the Vanderbilt University School of Engineering. MW acknowledges postdoctoral funding support from the Canadian Institute of Health Research (CIHR). LEP acknowledges a National Science Foundation Graduate Research Fellowship under grant number 1937963. Any opinions, findings, and conclusions or recommendations expressed in this material are those of the author(s) and do not necessarily reflect the views of the National Science Foundation.

