## Supplementary Information for "A Nanoparticle RIG-I Agonist for Cancer Immunotherapy"

### Supplementary Methods – DLin-MC3-DMA (MC3) Synthesis

<u>Overall Synthetic Scheme:</u>

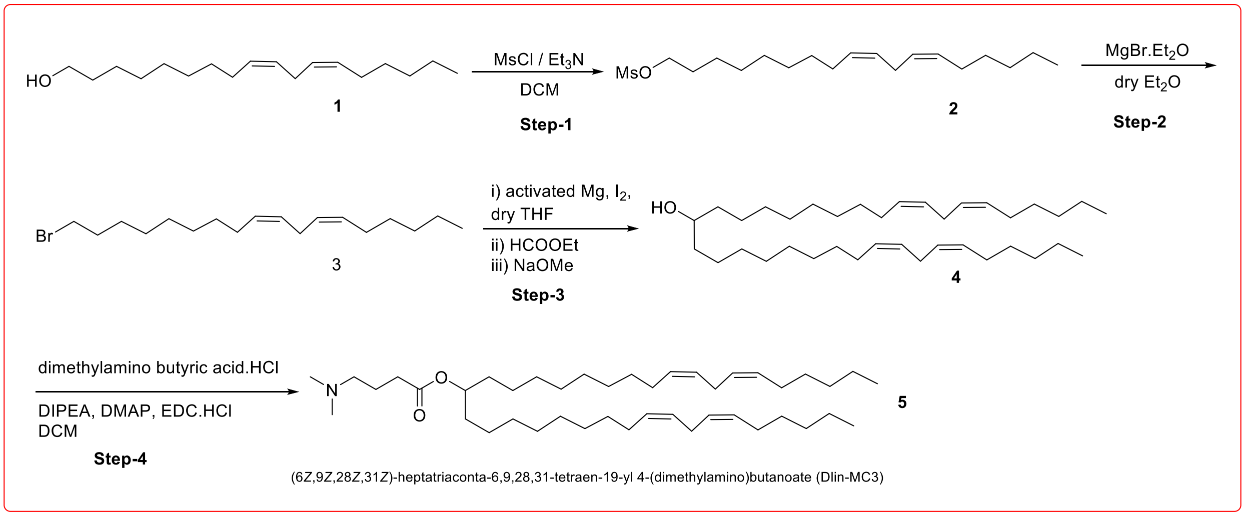


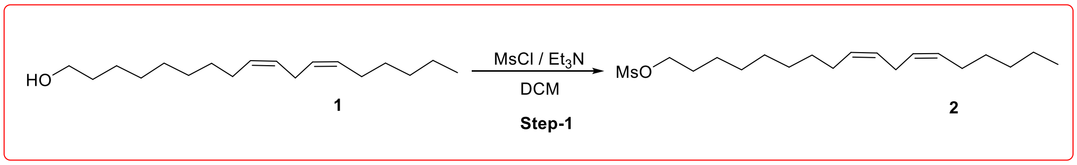


**Step-1:** (9Z,12Z)-octadeca-9,12-dien-1-yl methanesulfonate

Mesyl-Cl (0.702 ml, 9.01 mmol) was added dropwise to a stirred mixture of (9Z,12Z)-octadeca-9,12-dien-1-ol (2 g, 7.51 mmol) and TEA (4.29 ml, 30.77 mmol) in DCM (20 mL) at 0°C under argon. The resulting mixture was stirred at room temperature for 16 hours. The reaction mixture was diluted with DCM (50 mL) and washed with saturated aqueous NaCl (50 mL). The organic layer was dried over Na2SO4, filtered, and concentrated under reduced pressure to dryness to afford crude product. The resulting residue was purified by flash silica chromatography, elution gradient 0 to 100% EtOAc in hexanes. Product fractions were concentrated under reduced pressure to dryness to afford (9Z,12Z)-octadeca-9,12-dien-1-yl methanesulfonate (1.828 g, 70.7 %) as a colorless oil. $^{1}$H NMR (500MHz, CHLOROFORM-d) δ ppm 0.90 (t, *J* = 6.9 Hz, 3H), 1.24 - 1.47 (m, 16H), 1.69 - 1.82 (m, 2H), 2.06 (m, *J* = 7.3 Hz, 4H), 2.78 (t, 2H), 3.01 (s, 3H), 4.23 (t, *J* = 6.6 Hz, 2H), 5.27 - 5.46 (m, 4H).

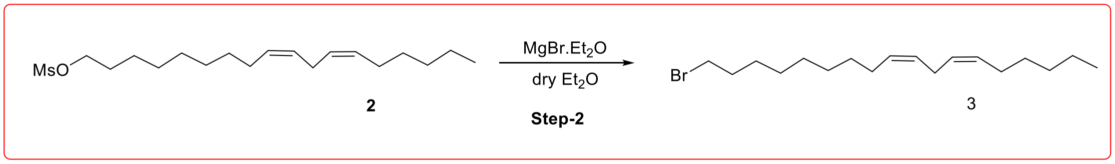


**Step-2:** (6Z,9Z)-18-bromooctadeca-6,9-diene

To a solution of (9Z,12Z)-octadeca-9,12-dien-1-yl methanesulfonate (1.828 g, 5.31 mmol) in THF (10 mL), lithium bromide (0.921 g, 10.61 mmol) was added in one portion under argon. The resulting clear colorless solution was heated to 65 °C and stirred for 16 hours. The reaction mixture was diluted with MTBE (30 mL) and washed with water (2 x 25 mL). The organic layer was dried (Na2SO4) and concentrated under reduced pressure to give crude product. The resulting residue was purified by flash silica chromatography, elution gradient 0 to 100% EtOAc in hexanes. Product fractions were concentrated under reduced pressure to dryness to afford (6Z,9Z)-18-bromooctadeca-6,9-diene (1.571 g, 90 %) as a colorless oil. 1H NMR (500MHz, CHLOROFORM-d) δ ppm 0.90 (t, J = 6.9 Hz, 3H), 1.24 - 1.49 (m, 16H), 1.87 (m, 2H), 2.06 (t, J = 7.0 Hz, 4H), 2.79 (s, 2H), 3.42 (t, J = 6.9 Hz, 2H), 5.26 - 5.48 (m, 4H).

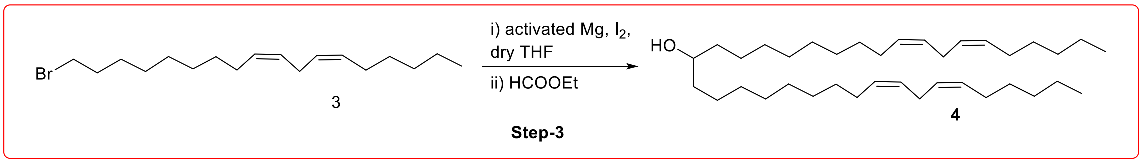


**Step-3:** (6Z,9Z,28Z,31Z)-heptatriaconta-6,9,28,31-tetraen-19-ol

Mg turnings (1.5 g, 0.0625 mole, 1.12 eq) and two crystals of iodine were taken in to clean dry 250 mL 2 neck round bottom flask. (6Z,9Z)-18-bromooctadeca-6,9-diene (1.34 g, 0.0558 mole, 1 eq) dissolved in 50 mL of THF was added slowly dropwise at room temperature. Exothermic conditions were observed. After completion of the addition, reaction mixture was heated at 50°C for 5h. The progress of the reaction was monitored by TLC. Reaction mixture was slowly allowed to come at 0°C and ethyl formate (4.26 g, 0.0575 mole, 1.03 eq) dissolved in 10 mL of THF was added drop wise at 0°C. Reaction mixture was slowly allowed to come at RT and continued for overnight with proper stirring, the reaction was monitored by TLC. The reaction mixture was then cooled to 0°C and quenched with 2 M HCl solution (100 mL). The reaction the mixture was diluted with ethyl acetate (100 mL) and separated. The aqueous layer was extracted with ethyl acetate (3 x 50 mL) and the combined organic layers were washed with brine solution (1 x 50 mL). The organics were dried over $Na_2SO_4$, filtered, and concentrated under reduced pressure to obtained light yellow oil. The compound was purified by flash column chromatography (silica gel, 0-10% Ethyl acetate in hexanes) and the pure product fractions were evaporated to afford (6Z,9Z,28Z,31Z)-heptatriaconta-6,9,28,31-tetraen-19-ol (1.4 g, 60%) as yellow oil. 1H NMR (500MHz, CHLOROFORM-d) δ ppm 0.87 – 0.91 (t, 6H), 1.20 - 1.43 (m, 46H), 2.01 – 2.06 (m, 9H), 2.75 – 2.79 (t, 4H), 3.58 (m, 1H), 5.29 – 5.43 (m, 8H).

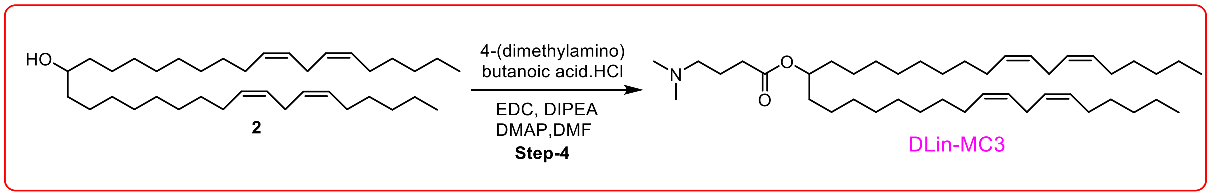


**Step-4:**(6Z,9Z,28Z,31Z)-heptatriaconta-6,9,28,31-tetraen-19-yl 4 (dimethylamino)butanoate (DLin-MC3-DMA)

Dimethylaminobutyric acid hydrochloride (0.950 g, 0.00567 mole, 2 eq) was added in one portion to (6Z,9Z,28Z,31Z)-heptatriaconta-6,9,28,31-tetraen-19-ol (1.5 g, 0.00284 mole, 1 eq) was dissolved in DMF (50 mL). The reaction mixture was cooled to 0°C and diisopropylethylamine (58 mL, 0.3308mole, 5 eq) and DMAP (1.61 g, 0.01323 mole, 0.2 eq) were added. After stirring for 5 min, EDC (2.37 g, 0.00567 mole, 2 eq) was added at 0°C The reaction was continued at room temperature for 16 h and monitored by TLC. The reaction mixture was cooled to 0°C and quenched with 10% citric acid solution (50 mL). The aqueous solution was extracted with DCM (3 x 50 mL) and the combined organic layers were washed with brine solution (50 mL). The organic layer was dried over $Na_2SO_4$, filtered, and concentrated under reduced pressure to obtained crude oil compound. The compound was purified by flash column chromatography (silica gel, 0-40% [20% MeOH and 1% NH4OH in DCM] in DCM). Product fractions were concentrated under reduced pressure to dryness to afford (6Z,9Z,28Z,31Z)-heptatriaconta-6,9,28,31-tetraen-19-yl 4-(dimethylamino)butanoate (1.42 g, 78 %) as a pale yellow oil. 1H NMR (500MHz, CHLOROFORM-d) δ ppm 0.89 (t, J = 6.8 Hz, 6H), 1.33 (m, 44H), 1.53 (d, 6.2 Hz, 4H), 2.04 (m, 11H), 2.53 (t, 6.4 Hz, 2H), 2.77 (t, J = 6.6 Hz, 4H), 2.80 (s, 2H), 3.00 (s, 6H), 3.26 (t, 7.7 Hz, 2H), 4.85 (m, 1H), 5.36 (m, 8H).

**Supplementary Data:**

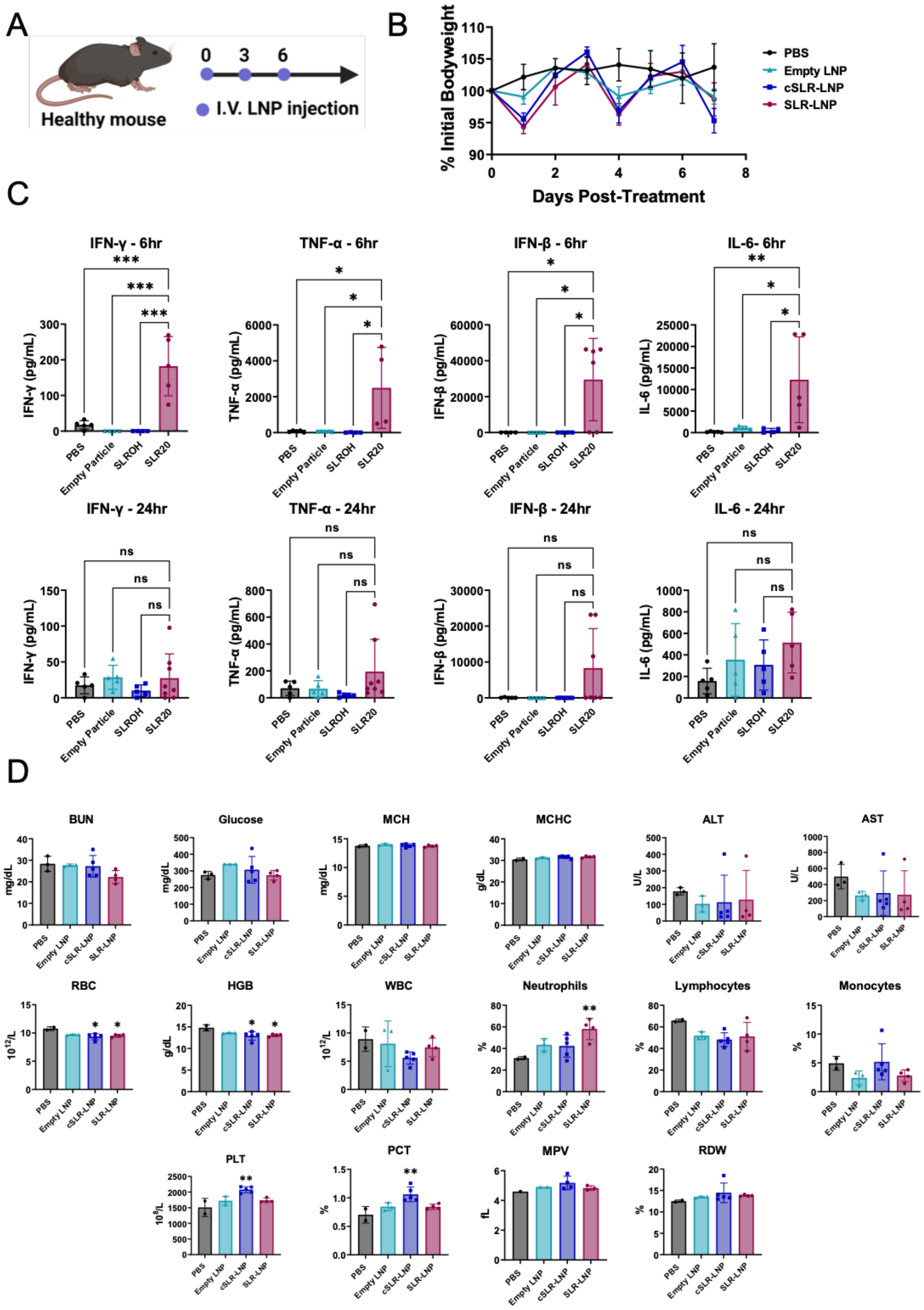


**Figure S1. Evaluation of toxicity of SLR-LNPs in mice. (A)** Schematic diagram showing the treatment schedule of healthy mice treated with 10 μg/mL SLR-LNP or control. **(B)** Body weight change of the mice after designated treatments. **(C)** Quantification of plasma cytokines 6 hours and 24 hours after the final treatment (n=3-5, *P ≤ 0.05, **P ≤ 0.01, ***P ≤ 0.001 by one-way ANOVA). **(D)** Blood biochemistry of healthy non-tumor-bearing syngeneic C57BL/6 mice. Animals (n=3-5) were administered 3 treatments and sacrificed 24 hours after final the final treatment. Blood samples were collected and analyzed to determine changes in RBCs, white blood cells (WBCs), neutrophils, platelets, and lymphocytes. Serum samples were used to analyze liver and kidney function, by measuring changes in ALT/AST, blood urea nitrogen (BUN), and creatinine. No significant changes were observed. (*P ≤ 0.05, **P ≤ 0.01 compared to PBS by one-way ANOVA).

1H NMR:

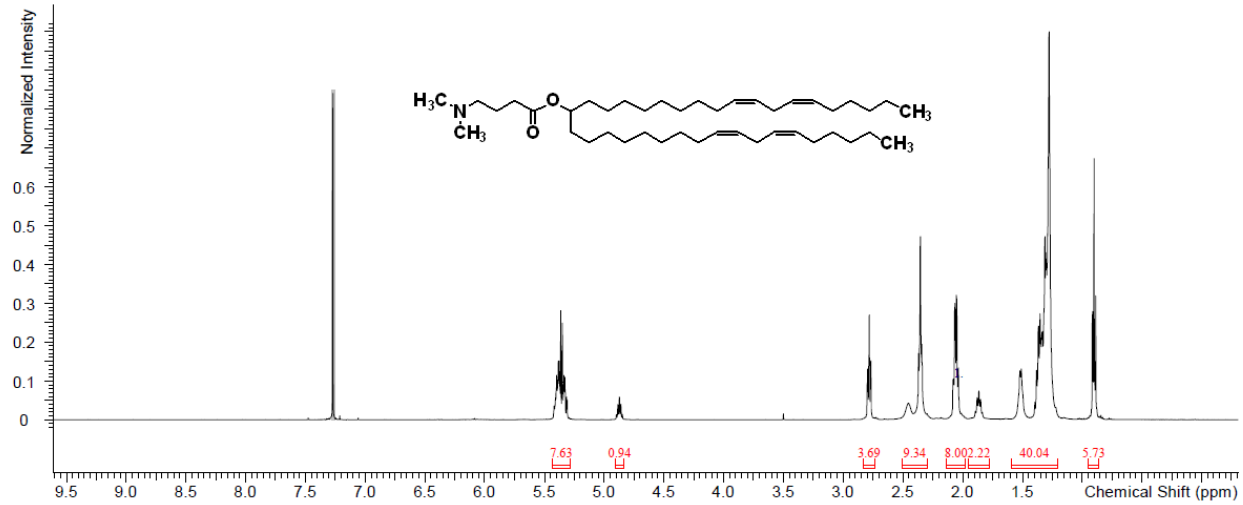


UPLC-ELSD:

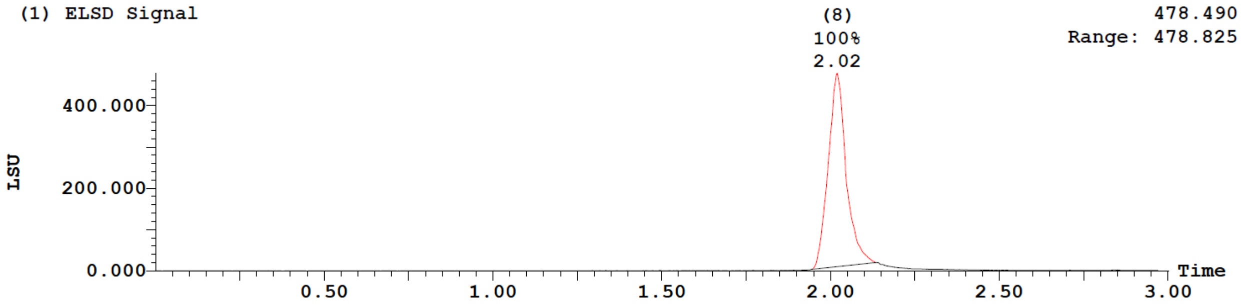


| Peak Number | Compound | Time | AreaAbs | Area %Total | Width | Height | Mass Found |
|---|---|---|---|---|---|---|---|
| 8 | | 2.02 | 3e+004 | 100.00 | 0 | 5e+005 | Not Found |

MS:

| Peak ID | Compound | Time | Mass Found |
|---|---|---|---|
| 8 | | 2.04 | Not Found |

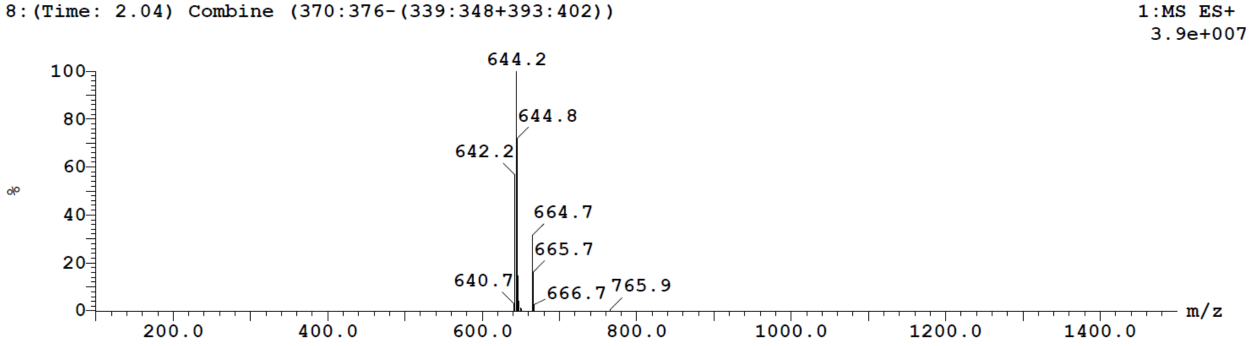


**Figure S2:** Chemical Characterization of DLin-MC3-DMA.

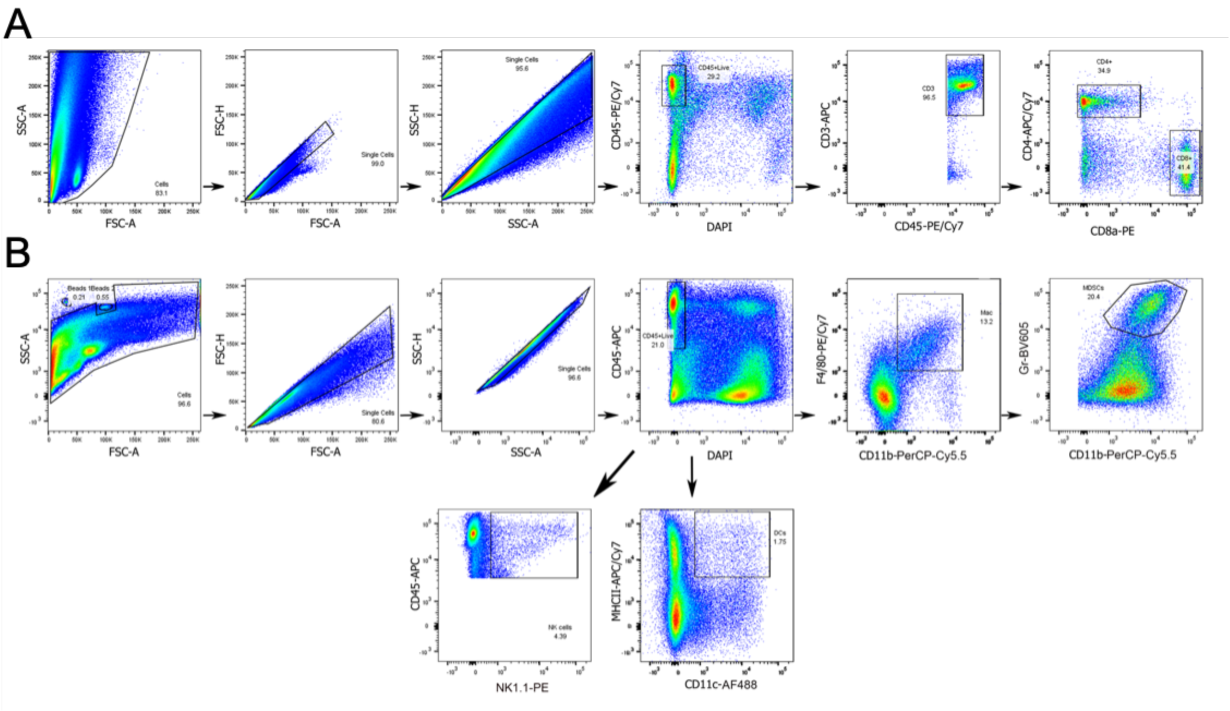


**Figure S3:** Representative flow cytometry dot plots showing gating strategy related to Figure 5B for analysis of tumor infiltrating **(A)** CD4$^+$ and CD8$^+$ T cells and **(B)** macrophages (CD11b$^+$F4/80$^+$), natural killer cells (NK 1.1$^+$), dendritic cells (CD11c$^+$MHCII$^+$), and MDSCs (CD11b$^+$Gr-1$^+$).
